# Molecular Evolutionary Analyses of Tooth Genes Support Sequential Loss of Enamel and Teeth in Baleen Whales (Mysticeti)

**DOI:** 10.1101/2021.11.10.468114

**Authors:** Jason G. Randall, John Gatesy, Mark S. Springer

## Abstract

The loss of teeth and evolution of baleen racks in Mysticeti was a profound transformation that permitted baleen whales to radiate and diversify into a previously underutilized ecological niche of bulk filter-feeding on zooplankton and other small prey. Ancestral state reconstructions suggest that teeth were lost in the common ancestor of crown Mysticeti. Genomic studies provide some support for this hypothesis and suggest that the genetic toolkit for enamel production was inactivated in the common ancestor of living baleen whales. However, molecular studies to date have not provided direct evidence for the complete loss of teeth, including their dentin component, on the stem mysticete branch. Given these results, several questions remain unanswered: (1) Were teeth lost in a single step or did enamel loss precede dentin loss? (2) Was enamel lost early or late on the stem mysticete branch? (3) If enamel and dentin/tooth loss were decoupled in the ancestry of baleen whales, did dentin loss occur on the stem mysticete branch or independently in different crown mysticete lineages? To address these outstanding questions, we compiled and analyzed complete protein-coding sequences for nine tooth-related genes from cetaceans with available genome data. Seven of these genes are associated with enamel formation (*ACP4, AMBN, AMELX, AMTN, ENAM, KLK4, MMP20*) whereas two other genes are either dentin-specific (*DSPP*) or tooth-specific (*ODAPH*) but not enamel-specific. Molecular evolutionary analyses indicate that all seven enamel-specific genes have inactivating mutations that are scattered across branches of the mysticete tree. Three of the enamel genes (*ACP4, KLK4, MMP20*) have inactivating mutations that are shared by all mysticetes. The two genes that are dentin-specific (*DSPP*) or tooth-specific (*ODAPH*) do not have any inactivating mutations that are shared by all mysticetes, but there are shared mutations in Balaenidae as well as in Plicogulae (Neobalaenidae + Balaenopteroidea). These shared mutations suggest that teeth were lost at most two times. Shared inactivating mutations and dN/dS analyses, in combination with cetacean divergence times, were used to estimate inactivation times of genes and by proxy enamel and tooth phenotypes. The results of these analyses are most compatible with a two-step model for the loss of teeth in the ancestry of living baleen whales: enamel was lost very early on the stem Mysticeti branch followed by the independent loss of dentin (and teeth) in the common ancestors of Balaenidae and Plicogulae, respectively. These results imply that some stem mysticetes, and even early crown mysticetes, may have had vestigial teeth comprised of dentin with no enamel. Our results also demonstrate that all odontocete species (in our study) with absent or degenerative enamel have inactivating mutations in one or more of their enamel genes.

## 1. Introduction

Cetaceans are a diverse group of fully aquatic mammals that exhibit a medley of morphological, physiological, and behavioral specializations that evolved in conjunction with their invasion and conquest of aquatic habitats. This clade includes both the largest known vertebrate (*Balaenoptera musculus* [blue whale]) and the mammal with the longest lifespan (*Balaena mysticetus* [bowhead whale]) (Gatesy et al., 2013; Keane et al., 2015). Aside from their remarkable phenotypic diversity, ceteceans have an incredible and well-documented macroevolutionary history that is illuminated by both fossils and genomes (Bajpai et al., 2009; Thewissen et al., 2009; Gatesy et al., 2013; McGowen et al., 2014, 2020b; Berta et al., 2016). Recent decades of paleontological research have yielded extinct species that document the transition from life on land to the aquatic realm (Thewissen and Bajpai, 2001; Thewissen et al., 2001, 2009; Gingerich, 2012; Gatesy et al., 2013). These fossils record the evolution of key morphological features of the cetacean body plan including development of paddle-shaped forelimbs, ‘telescoping’ of the skull, and reduction of the hindlimbs (Bejder and Hall, 2002; Gatesy et al., 2013). At the genomic level, molecular evolutionary studies provide evidence for both positive selection and extensive gene loss in association with the transition from land to water. These changes are linked to a variety of anatomical structures and organ systems that were modified in the ancestor of Cetacea including the skin, limbs, lungs, pineal gland, brain, brown fat, eyes, ears, vomeronasal organ, and nose (Gatesy et al., 2013; Nery et al., 2013; McGowen et al., 2014, 2020b; Gaudry et al., 2017; Sharma et al., 2018; Huelsmann et al., 2019; Thermudo et al., 2020; Emerling et al., 2021; Springer and Gatesy, 2018; Springer et al., 2021).

During the late Eocene (∼36-37 million years ago [Ma]) there was a cladogenic split in the last common ancestor of Neoceti (crown Cetacea) that resulted in the reciprocally monophyletic clades Odontoceti (toothed whales) and Mysticeti (baleen whales) (McGowen et al., 2009, 2020a; Gatesy et al. 2013). Whereas most odontocetes still possess enamel-capped teeth like their terrestrial ancestors, extant mysticetes have lost their teeth and instead feed using baleen racks that consist of keratin (Uhen, 2010; Gatesy et al., 2013). The evolution of baleen was associated with a profound dietary transformation that allowed mysticetes to exploit a previously underutilized food resource, zooplankton and other tiny prey, by bulk filter-feeding instead of raptorial or suction feeding on individual prey as in most stem cetaceans and odontocetes.

Diverse data support the hypothesis that baleen whales evolved from toothed ancestors: (1) cladistic analyses of phenotypic data and ancestral state reconstructions document the evolution of edentulous mysticetes from stem mysticete ancestors that possessed teeth (Fitzgerald, 2006, 2009; Uhen, 2010; Meredith et al., 2011; Gatesy et al., 2013), (2) the presence of molecular relics of both enamel-related and dentin/tooth-related tooth genes in the genomes of extant mysticetes (Deméré et al., 2008; Meredith et al., 2009, 2011a; Kawasaki et al., 2014; Berta et al., 2016; Springer et al., 2016a, 2019; Gatesy et al., 2021; Mu et al., 2021), and (3) the initiation of tooth formation in whale fetuses (i.e., tooth germs that are sometimes mineralized and resorbed before birth) (Dissel-Scherft and Vervoort, 1954; Thewissen and Williams, 2002; Deméré et al., 2008; Thewissen et al., 2017; Lanzetti, 2019). Note that our use of ‘teeth’ in the remainder of this paper refers to postnatal teeth and does not refer to tooth germs that occur in mysticete fetuses.

However, what has been less clear is how and when postnatal teeth were lost, and if tooth loss occurred before or after the evolution of baleen. The co-occurrence hypothesis suggests that baleen evolved before tooth loss and that teeth and baleen functioned together before teeth were subsequently lost in the ancestry of extant mysticetes (Deméré et al., 2008; Boessenecker and Fordyce, 2015; Ekdale and Deméré, 2021). A variation of this hypothesis is the dentral filtration hypothesis, which contends that early mysticetes used their teeth to filter feed before developing baleen for this purpose before eventually losing their teeth (Geisler et al., 2017; also see Hocking et al. [2017] for opposing view). By contrast with these two hypotheses wherein baleen evolved before functional teeth were lost, the toothless suction-feeding hypothesis postulates that stem mysticetes lost their teeth and were edentulous suction feeders prior to the evolution of baleen (Peredo et al., 2017, 2018).

Deméré et al. (2008) advocated for the co-occurrence hypothesis based on the presence of both teeth and inferred baleen (medial to the teeth) in *Aetiocetus weltoni* and two additional Oligocene mysticete species. Baleen was hypothesized based on the presence of lateral palatal foramina and sulci that are used to supply blood and nerves to the ever-growing baleen racks in extant baleen whales (Deméré et al., 2008). If the co-occurrence hypothesis is correct, the lateral palatal foramina in *A. weltoni* should connect internally to the superior alveolar canal, which transmits blood vessels and nerves to the maxillary teeth in odontocetes and to the baleen racks in extant mysticetes (Edkale and Deméré, 2021). Ekdale and Deméré (2021) used CT scan images of *A. weltoni* to show that this early mysticete exhibits a condition that is intermediate between a representative odontocete (*Tursiops*) and a representative mysticete (*Eschrichtius*). Specifically, the lateral branch of the maxillary canal (superior alveolar canal) has connections with both the dental alveoli and the lateral palatal foramina. Ekdale and Deméré’s (2021) results provide evidence for the co-occurrence hypothesis, but do not bear on the timing of enamel versus dentin loss in the ancestor of modern baleen whales.

To clarify the correct sequence of evolutionary events that culminated in tooth loss, several studies have investigated the functional versus pseudogenic status of tooth-specific genes in extant mysticetes. If teeth were lost on the stem mysticete branch (Fitzgerald, 2006, 2009; Meredith et al., 2011a), we should expect to find inactivating mutations in both enamel and dentin related genes that are shared by all extant mysticetes (Deméré et al., 2008; Meredith et al., 2009, 2011a; Berta et al., 2016; Springer et al., 2016a, 2019). Inactivating mutations include genetic changes that are expected to radically impact or impair a gene’s function including frameshift insertions and deletions [frameshift indels], altered start or stop codons, premature stop codons, insertion of long retroelement sequences, modified splice sites at intron/exon boundaries, and deletions of an exon(s) or an entire gene. Meredith et al. (2011a) reported the first evidence of a shared inactivating mutation in mysticetes. Specifically, the enamel-specific gene matrix metallopeptidase 20 (*MMP20*) has a shared retroelement insertion (a CHR-SINE in exon 2 of all living mysticetes that were examined. More recently, Mu et al. (2021) and Gatesy et al. (2021) reported inactivating mutations in acid phosphatase 4 (*ACP4*) and kallikrein related peptidase 4 (*KLK4*), respectively, that are shared by all extant mysticetes that were examined. Like *MMP20*, these genes are related to enamel formation. Other enamel-specific genes (amelogenin X-linked [*AMELX*], ameloblastin [*AMBN*], amelotin [*AMTN*], enamelin [*ENAM*]) have inactivating mutations in multiple mysticetes, but none that are shared by all mysticete species for the exons that were investigated (Deméré et al., 2008; Meredith et al., 2009, 2011a; Gatesy et al., 2021). The absence of shared inactivating mutations in these enamel-specific genes may reflect a lag time between the initial relaxation of selection pressures for the maintenance of enamel on the stem mysticete branch and the first occurrence of an inactivating mutation in an enamel gene. Given the decelerated rate of molecular evolution in cetaceans relative to most other mammals, such mutational lags are predicted (Meredith et al., 2009).

Two other genes are dentin-specific (dentin sialophosphoprotein [*DSPP*]) (McKnight and Fisher, 2009) or tooth-specific (odontogenesis associated phosphoprotein [*ODAPH*]) (Springer et al., 2016a), but not enamel-specific. DSPP plays an essential role in the formation of dentin and the protease-processed products of DSPP comprise the largest component of non-collagenous proteins found in dentin (Yamakoshi and Simmer, 2018). Mutations in the human *DSPP* gene are known to cause both dentin dysplasia and dentinogenesis imperfecta (Yamakoshi and Simmer, 2018). *DSPP* exhibits a 1-bp frameshift insertion that is shared by two balaenids (*Balaena mysticetus, Eubalaena japonica*) (Gatesy et al., 2021). However, this exon is deleted in three balaenopteroids that were examined, and it is unclear whether the 1-bp deletion in exon 3 occurred in the ancestor of balaenids or instead is an older mutation that occurred in the common ancestor of extant mysticetes that was subsequently erased in balaenopteroids by the much larger deletion that completely removed exon 3. If the balaenid 1-bp deletion is also present in the neobalaenid *Caperea marginata*, which is the sister group of balaenopteroids, this would provide evidence for inactivation of the genetic toolkit for dentin production on the stem lineage to crown Mysticeti.

In the case of *ODAPH*, this gene is intact in all dentate placental mammals that have been investigated including several species with enamelless teeth (*Orycteropus afer* [aardvark], *Dasypus novemcinctus* [nine-banded armadillo], *Tolypeutes matacus* [southern three-banded armadillo], *Chaetophractus vellerosus* [screaming hairy armadillo], *Choloepus hoffmanni* [Hoffmann’s two-toed sloth], *Choloepus didactylus* [Linnaeus’s two-toed sloth]) (Springer et al., 2016; Gatesy et al., 2021). Morever, Springer et al. (2016a) found that dN/dS values on branches leading to species with enamelless teeth are not significantly different from dN/dS values on branches leading to species with enamel-capped teeth. These results suggest that *ODAPH* remains under purifying selection even in species that have lost the enamel caps on their teeth. By contrast, all toothless placental mammals that have been investigated (ten mysticetes, *Manis pentadactyla* [Chinese pangolin], *Tamandua tetradactyla* [collared anteater]) have inactivating mutations in *ODAPH* including complete deletion of this locus in most balaenopteroids (Springer et al., 2016; Gatesy et al., 2021). The inactivation of *ODAPH* in multiple toothless clades, but not enamelless clades, suggests that *ODAPH* is tooth-specific but not enamel-specific. At the same time, *ODAPH* has a role in enamel formation and mutations in this gene are known to cause amelogenesis imperfecta (Parry et al., 2012; Ji et al., 2021; Liang et al., 2021). Even though all mysticetes that have been investigated have inactivating mutations in *ODAPH*, there are no inactivating mutations in the protein-coding sequences of this gene that are shared by all mysticetes. However, this gene was lost at most three times based on inactivating mutations that are shared by Balaenidae and by Balaenopteroidea, respectively; the *Caperea* genome has not been examined to date and could document a third independent inactivation of this gene (Springer et al., 2016a). Importantly, the absence of inactivating mutations shared by all extant mysticetes in *DSPP* and in *ODAPH* does not imply that these genes were necessarily functional in the most recent common ancestor of extant baleen whales. As noted above, mysticetes have very slow rates of molecular evolution, and there may have been a lag time between relaxed selection on the dentition and the first occurrence of an inactivating mutation in one or both of these genes. Here, dN/dS analyses that measure selective constraints and gene inactivation dating have not yet been employed to examine the timing of relaxed selection on *DSPP* and *ODAPH*.

In view of the above several key questions remain unanswered: (1) Were teeth lost in a single step or did enamel loss precede dentin loss? (2) Was enamel lost early or late on the stem mysticete branch? (3) If enamel and tooth loss were decoupled in the ancestry of baleen whales, did tooth loss occur on the stem mysticete branch or independently in multiple crown mysticete lineages? To address these questions, we compiled and analyzed complete protein-coding sequences for nine tooth-specific genes from cetaceans with assembled genomes or raw Illumina data. Seven of these genes are enamel specific (*ACP4, AMBN, AMELX, AMTN, ENAM, KLK4, MMP20*) and two are related to dentin/tooth formation (*DSPP, ODAPH*) with the caveat that *ODAPH* is pleiotropic and is also important for enamel maturation (Liang et al., 2021). Taxon sampling included representatives of all four crown mysticete families (Balaenidae, Neobalaenidae, Eschrichtiidae, Balaenopteridae); a genomic library for *Caperea marginata* (Neobalaenidae) was used to sequence short reads to recover and assemble tooth gene sequences from this species. The nine genes listed above were investigated for shared inactivating mutations. We also employed dN/dS values to determine the timing of inactivation for both enamel and dentin genes, and by proxy the inferred loss of both enamel and teeth.

## 2. Methods

### 2.1. Gene sampling

Nine genes were chosen for study based on prior evidence that these loci are enamel-specific or tooth-specific with respect to their essential functions that are maintained by natural selection (Deméré et al., 2008; McKnight and Fisher, 2009; Meredith et al., 2009; 2011a, 2013, 2014; Gasse et al., 2012; Kawasaki et al., 2014; Springer et al., 2015, 2016a, 2019; Mu et al., 2021; Gatesy et al., 2021). Each of these nine genes has become pseudogenized in one or more clades of enamelless or edentulous (toothless) mammals. A tenth tooth-related gene (*ODAM*) is also inactivated in enamelless and edentulous mammals, but this gene is also pseudogenized in all toothed whales that were investigated as well as several other clades of mammals with enamel-capped teeth (Springer et al., 2019). For this reason we omitted *ODAM* from our study. Also, exon 4 of *AMELX* was not included in our analyses because this exon is subject to alternative splicing and is absent in many mammals (Delgado et al., 2005; Sire et al., 2005, 2006, 2007).

### 2.2. Taxon sampling

Taxon sampling for this study included 44 species of which 13 were mysticetes, 14 were odontocetes, and 17 were terrestrial or semiaquatic cetartiodactyl outgroups. Mysticetes included *Balaena mysticetus* (bowhead whale), *Balaenoptera acutorostrata* (common minke whale), *Balaenoptera bonaerensis* (Antarctic minke whale), *Balaenoptera borealis* (sei whale), *Balaenoptera edeni* (Bryde’s whale) *Balaenoptera musculus* (blue whale), *Balaenoptera physalus* (fin whale), *Caperea marginata* (pygmy right whale), *Eschrichtus robustus* (gray whale), *Eubalaena glacialis* (North Atlantic right whale), *Eubalaena australis* (southern right whale), *Eubalaena japonica* (North Pacific right whale), and *Megaptera novaeangliae* (humpback whale). Odontocetes included *Delphinapterus leucas* (beluga), *Kogia breviceps* (pygmy sperm whale), *Kogia sima* (dwarf sperm whale), *Lagenorhynchus obliquidens* (Pacific white-sided dolphin), *Lipotes vexillifer* (Chinese river dolphin), *Mesoplodon bidens* (Sowerby’s beaked whale), *Monodon monoceros* (narwhal), *Neophocaena asiaeorientalis* (Yangtze finless porpoise), *Orcinus orca* (killer whale), *Physeter macrocephalus* (sperm whale), *Phocoena phocoena* (harbor porpoise), *Sousa chinensis* (Indo-Pacific humpbacked dolphin), *Tursiops aduncus* (Indo-Pacific bottlenose dolphin), and *Tursiops truncatus* (common bottlenose dolphin). Outgroup taxa included *Bison bison* (American bison), *Bos mutus* (wild yak), *Bubalus bubalis* (water buffalo), *Camelus bactrianus* (Bactrian camel), *Capra hircus* (domestic goat), *Catagonus wagneri* (Chacoan peccary), *Choeropsis liberiensis* (pygmy hippopotamus), *Elaphurus davidianus* (Pere David’s deer), *Giraffa camelopardalis* (giraffe), *Hippopotamus amphibius* (river hippopotamus), *Moschus moschiferus* (Siberian musk deer), *Odocoileus virginianus* (white-tailed deer), *Okapia johnstoni* (okapi), *Ovis aries* (domestic sheep), *Sus scrofa* (wild boar), *Tragulus javanicus* (Java mouse-deer), and *Vicugna pacos* (alpaca).

### 2.3. Data collection

DNA sequences for nine different tooth genes (*ACP4, AMBN, AMELX, AMTN, DSPP, ENAM, KLK4, MMP20, ODAPH*) were obtained from (1) assembled genomes at NCBI (https://www.ncbi.nlm.nih.gov/) and the The Bowhead Whale Genome Resource (http://www.bowhead-whale.org/), (2) raw sequence reads at NCBI’s Sequence Read Archive (SRA), and (3) newly generated Illumina whole-genome sequence data (J.G, M.S.S.) (Table S1). NCBI’s RefSeq and Nucleotide databases were searched using keywords for all nine genes in conjunction with taxon names for four reference species (*Capra hircus, Camelus bactrianus, Orcinus orca, Tursiops truncatus*). Sequences for each reference species were then imported into Geneious Prime (current version 2021.1.1, https://geneious.com) (Kearse et al., 2012), aligned with MAFFT (Katoh and Toh, 2008), and cross-checked against each other for consistent annotations. Sequences for additional species were collected through NCBI’s Nucleotide Basic Local Alignment Search Tool (BLAST), which was used to search both assembled and unassembled genomes using the whole-genome shotgun (WGS) and SRA databases, respectively. Each BLAST search employed a query sequence from a closely related species. Megablast was used for highly similar sequences (e.g., taxa in same family), whereas Blastn was used for less similar sequences (e.g., taxa in different families). Top-scoring BLAST results were imported into Geneious Prime. Sequences obtained through the SRA database were assembled using Geneious Prime’s ‘Map to Reference’ approach, where the reference sequence was a closely related species to the SRA taxon. We allowed for a maximum mismatch of 10% per read and required a minimum of two reads for base calling with a consensus threshold of 65%. Unassembled genome sequences for two additional cetacean species (*Caperea marginata, Kogia sima*) were obtained from DNA libraries that were constructed with Illumina’s NeoPrep procedure and then sequenced at ∼40X coverage at the New York Genome Center with paired-end sequencing (150 bp per read) on a HiSeq 2500 platform. DNA samples for these libraries were provided by Southwest Fisheries Science Center (SWFSC) (*C. marginata* [Lab ID 5989]; *K. sima* [Lab ID 175303]). Sequences for the nine tooth genes were then obtained using a map to reference approach as described above. Accession numbers for these new sequences are OK282856-OK282863 for *C. marginata* and OK391138-OK391144 for *K. sima* (Table S1).

### 2.4. Alignments and tabulation of inactivating mutations

Complete protein-coding sequences and introns were aligned in Geneious Prime using MAFFT (Katoh and Toh, 2008). Sequences were manually spot-checked for alignment errors using AliView version 1.23 (Larsson, 2014). Alignments were examined for inactivating mutations (frameshift indels, start and stop codon mutations, premature stop codons, splice site mutations), which were annotated in Geneious Prime. Mutations were mapped on to the species tree using delayed transformation (DELTRAN) parsimony optimization.

### 2.5. Phylogenetic analyses

Gene trees were constructed from complete protein-coding sequences with maximum likelihood using the program RAxML version 8.2.11 in Geneious Prime (raxmlHPC-SSE3-MAC) (Stamatakis, 2014). Rapid bootstrapping (500 pseudoreplicates) and a search for the best tree were performed in the same analysis (Stamatakis et al., 2008). We employed the GTRGAMMA option, which implements the GTR + Γ model of sequence evolution.

### 2.6. Selection analyses

Inferred inactivating mutations, estimates of selection intensity (dN/dS analyses), and divergence times from McGowen et al.’s (2020a) cetacean timetree were used to reconstruct inactivation times of tooth genes and, by proxy, phenotypes. Selection (dN/dS) analyses were conducted with the codeml program of PAML (version 4.9e; Yang, 2007). Analyses were performed with a concatenation of seven enamel-specific genes (*ACP4, AMBN, AMELX, AMTN, ENAM, KLK4, MMP20*) that serve as a proxy for enamel and two dentin/tooth-specific genes (*DSPP, ODAPH*) that serve as a proxy for dentin/teeth. We used concatenations of enamel and dentin/tooth genes rather than individual genes because larger data sets yield dN/dS that are less impacted by sampling error. For the analysis with *DSPP* + *ODAPH* we excluded exon 4 of *DSPP* because this exon contains short, highly repetitive motifs that are difficult to align. We employed branch-specific codon models with branch categories (background, transitional, pseudogenic) that were based on phenotypes and shared genetic mutations (*sensu* Meredith et al., 2009). The background branch category includes branches that lead to internal nodes or extant species with functional teeth that are capped with prismatic enamel (seven enamel genes) or teeth irrespective of whether enamel is present (two dentin/tooth genes). These branches are expected to have evolved under purifying selection with dN/dS < 1 in terrestrial cetartiodactyl outgroups and some or all odontocetes (all odontocetes for dentin/tooth genes but only some odontocetes for enamel genes). Transitional branches lead to internal nodes or extant species that lack enamel/prismatic enamel or teeth and contain the first detected occurrence of an inactivating mutation that is shared by all members of a clade. Each transitional branch was given its own branch category. Transitional branches have mixed evolutionary histories that include a period of evolution under purifying selection followed by a period of neutral evolution after selection was relaxed and the phenotype was lost (Meredith et al., 2009). DN/dS values on transitional branches are expected to be intermediate between dN/dS values for background branches and pseudogenic branches. Pseudogenic branches post-date transitional branches and are expected to have neutral evolutionary histories with dN/dS values near 1. DN/dS analyses for the seven enamel genes included nine branch categories (two background [#0, #1], six transitional [#2-#7], one pseudogenic [#8]) as follows: #0 background terrestrial cetartiodactyl outgroups, #1 background Odontoceti (all odontocetes with complex enamel [categories 4 and 5 of Werth et al., 2020]), #2 *Physeter*, #3 stem *Kogia*, #4 *Mesoplodon*, #5 stem Monodontidae, #6 stem Phocoenidae, #7 stem Mysticeti, #8 crown Mysticeti + crown *Kogia* + crown Monodontidae + crown Phocoenidae. Background cetartiodactyl outgroups and background Odontoceti were separated into two categories to determine if there is relaxed selection on these enamel genes in odontocetes with complex enamel, i.e., enamel with prismatic radial enamel that may also exhibit Hunter-Schreger bands (HSB) or other decussations (Werth et al., 2020). DN/dS analyses with the dentin genes included three branch categories: #0 background terrestrial cetartiodactyl outgroups plus stem and crown Odontoceti, #1 stem Mysticeti + stem Balaenidae, #2 crown Balaenidae. Other mysticete families were excluded from these analyses because the coding sequences for exons 1-3 of *DSPP* and all of *ODAPH* have been deleted (*Balaenoptera musculus* retains *ODAPH* [Springer et al., 2016a] but not exons 1-3 of *DSPP*).

Analyses were performed with two codon frequency models, CF1 and CF2 (Yang, 2007). CF1 estimates codon frequencies from mean nucleotide frequencies across all three codon positions, whereas CF2 estimates frequencies at each of the individual codon positions. Analyses were conducted with both fixed (dN/dS = 1) and estimated values for the fully pseudogenic branch category (dN/dS = 1.0 or dN/dS = estimated); chi-square tests were conducted using the program chi2 (Yang, 2007) to determine whether the analyses with fixed versus estimated dN/dS values for the pseudogenic branch category were significantly different from each other. All frameshift insertions were deleted prior to performing dN/dS analyses. In addition, premature stop codons were recoded as missing data as required for codeml analyses. The species tree used for Cetartiodactyla relationships was taken from McGowen et al. (2020a) for cetaceans and Hassanin et al. (2012) for outgroups to Cetacea.

### 2.7. Gene inactivation times

Inactivation times for the concatenation of seven enamel genes and for the concatenation of two dentin genes were each estimated using equations from Meredith et al. (2009) that allow for either one or two synonymous substitution rates. The one synonymous substitution rate model assumes that the rate of synonymous substitution is neutral and equal on both functional and pseudogenic branches, whereas the two-rate model assumes that the synonymous substitution rate on functional branches is non-neutral and is 70% of the substitution rate on pseudogenic branches (Bustamante et al., 2002; Meredith et al., 2009). Divergence time estimates were taken from McGowen et al. (2020a).

### 2.8. Data availability

Alignments for protein-coding sequences for individual genes, gene trees, concatenated alignments for codeml analyses (frameshift insertions removed, stop codons replaced with Ns), species trees for codeml analyses, codeml outfiles, and an Excel calculator for gene inactivation times are posted at: https://figshare.com/s/411a3a1b8fe230ed9d85 [THIS IS A PRIVATE LINK THAT WILL BE REPLACED WITH A PERMANENT DOI IF THE MANUSCRIPT IS ACCEPTED].

## 3. Results

### 3.1. Alignments and gene trees

Complete coding sequences for all genes were recovered for all the non-cetacean cetartiodactyl outgroup taxa that were sampled, as well as for all odontocete species except for the dwarf and pygmy sperm whales (*Kogia* spp.) that lack well-developed enamel (Bianucci and Landini, 2006; Werth et al., 2020). Mysticetes that were sampled had varying degrees of completeness for the coding sequences of the nine tooth genes (*ACP4, AMBN, AMELX, AMTN, DSPP, ENAM, KLK4, MMP20, ODAPH*). Alignments for the coding sequences of each gene are provided in Supplementary Information. RAxML gene trees for each gene are provided in Figure S1. Table 1 summarizes the presence or absence of 27 well-supported and non-controversial clades in Cetartiodactyla (Meredith et al., 2011b, Hassanin et al., 2012; Gatesy et al., 2013; McGowen et al., 2014, 2020a) on the individual gene trees. All of the individual gene trees recovered the majority of well-supported clades in Cetartiodactyla. Eight clades were recovered on all gene trees with the constituent taxa and the mean number of recovered clades was 23.78. Coding sequences for *ENAM* are longer than for any of the other enamel-related genes and 27 of 27 clades were recovered on the *ENAM* tree. *ODAPH* contains the shortest coding sequences and recovered the fewest clades (20 of 25). Twenty-six of 27 clades were recovered on the *MMP20* gene tree. Gene trees for the other six genes recovered 22 to 25 of the well-supported clades.

**Table 1.** Monophyly of well-supported cetartiodactyl clades on maximum likelihood phylograms for nine tooth related genes.

| Clade | Gene |  |  |  |  |  |  |  |  |
| --- | --- | --- | --- | --- | --- | --- | --- | --- | --- |
|  | *ACP4* | *AMBN* | *AMELX* | *AMTN* | *DSPP* | *ENAM* | *KLK4* | *MMP20* | *ODAPH* |
| Camelidae | Yes | Yes | Yes | Yes | Yes | Yes | Yes | Yes | Yes |
| Suiformes | Yes | Yes | Yes | Yes | Yes | Yes | Yes | Yes | Yes |
| Ruminantia + Cetancodonta | Yes | Yes | Yes | Yes | Yes | Yes | Yes | Yes | No |
| Ruminantia | Yes | Yes | Yes | Yes | Yes | Yes | Yes | Yes | Yes |
| Pecora | Yes | Yes | Yes | Yes | Yes | Yes | Yes | Yes | Yes |
| Bovidae | Yes | No | Yes | Yes | Yes | Yes | Yes | Yes | No |
| Bovinae | Yes | Yes | Yes | No | Yes | Yes | Yes | Yes | Yes |
| Caprinae | Yes | Yes | Yes | Yes | Yes | Yes | Yes | Yes | Yes |
| Cervidae | Yes | Yes | Yes | Yes | Yes | Yes | Yes | Yes | Yes |
| Giraffidae | Yes | Yes | Yes | Yes | Yes | Yes | Yes | Yes | Yes |
| Cetancodonta | Yes | No | Yes | Yes | Yes | Yes | No | Yes | Yes |
| Hippopotamidae | Yes | Yes | Yes | Yes | Yes | Yes | Yes | Yes | Yes |
| Cetacea | Yes | Yes | Yes | Yes | Yes | Yes | Yes | Yes | Yes |
| Odontoceti | No | Yes | Yes | No | Yes | Yes | Yes | Yes | No |
| Physeteroidea | No | Yes | Yes | Yes | Yes | Yes | NA | Yes | No |
| *Kogia* | Yes | NA | Yes | Yes | Yes | Yes | NA | Yes | Yes |
| Delphinida + Ziphiidae | Yes | Yes | No | Yes | Yes | Yes | Yes | Yes | Yes |
| Delphinida | Yes | Yes | Yes | Yes | Yes | Yes | No | Yes | Yes |
| Delphinoidea | Yes | Yes | Yes | Yes | Yes | Yes | Yes | Yes | Yes |
| Monodontidae + Phocoenidae | Yes | Yes | No | Yes | No | Yes | Yes | Yes | Yes |
| Monodontidae | Yes | Yes | No | Yes | Yes | Yes | No | Yes | No |
| Phocoenidae | Yes | Yes | Yes | Yes | Yes | Yes | Yes | Yes | Yes |
| Delphinidae | Yes | Yes | Yes | Yes | Yes | Yes | Yes | No | Yes |
| Mysticeti | Yes | Yes | Yes | No | Yes | Yes | Yes | Yes | Yes |
| Balaenidae | Yes | Yes | Yes | Yes | Yes | Yes | Yes | Yes | Yes |
| Balaenopteroidea | No | No | Yes | Yes | Yes | Yes | Yes | Yes | NA |
| *Balaenoptera acutorostrata* + *B. bonaerensis* | Yes | No | Yes | Yes | No | Yes | Yes | Yes | NA |
| Total number of clades on gene tree | 24/27 | 22/26 | 24/27 | 24/27 | 25/27 | 27/27 | 22/25 | 26/27 | 20/25 |

### 3.2. Inactivating mutations in mysticetes

Table 2 provides a summary of inactivating mutations in mysticetes, which are also mapped onto a species tree for Mysticeti in Figure 1. Of the seven enamel genes, three (*ACP4, KLK4, MMP20*) were found to have inactivating mutations that are shared by all mysticetes. *ACP4* has a 1-bp deletion in exon 4; *KLK4* has a 1-bp deletion in exon 3; and *MMP20* has a SINE insertion in exon 2. Remnants of this insertion range from 302 to 324 bp in different mysticete species with assembled genomes, and all possible reading frames of this SINE insertion contain premature stop codons that would result in a severely truncated MMP20 protein (Meredith et al., 2011a). Mu et al. (2021) reported a second 1-bp deletion in exon 5 of *ACP4* that is shared by all mysticetes that were sampled in that study, but expanded taxon sampling revealed that this deletion is not shared by *Caperea marginata* or *Balaenoptera edeni*. Nevertheless, three shared inactivating mutations support the hypothesis that baleen whales lost functional enamel on the stem branch leading to crown Mysticeti (Fig. 1).

**Table 2.**
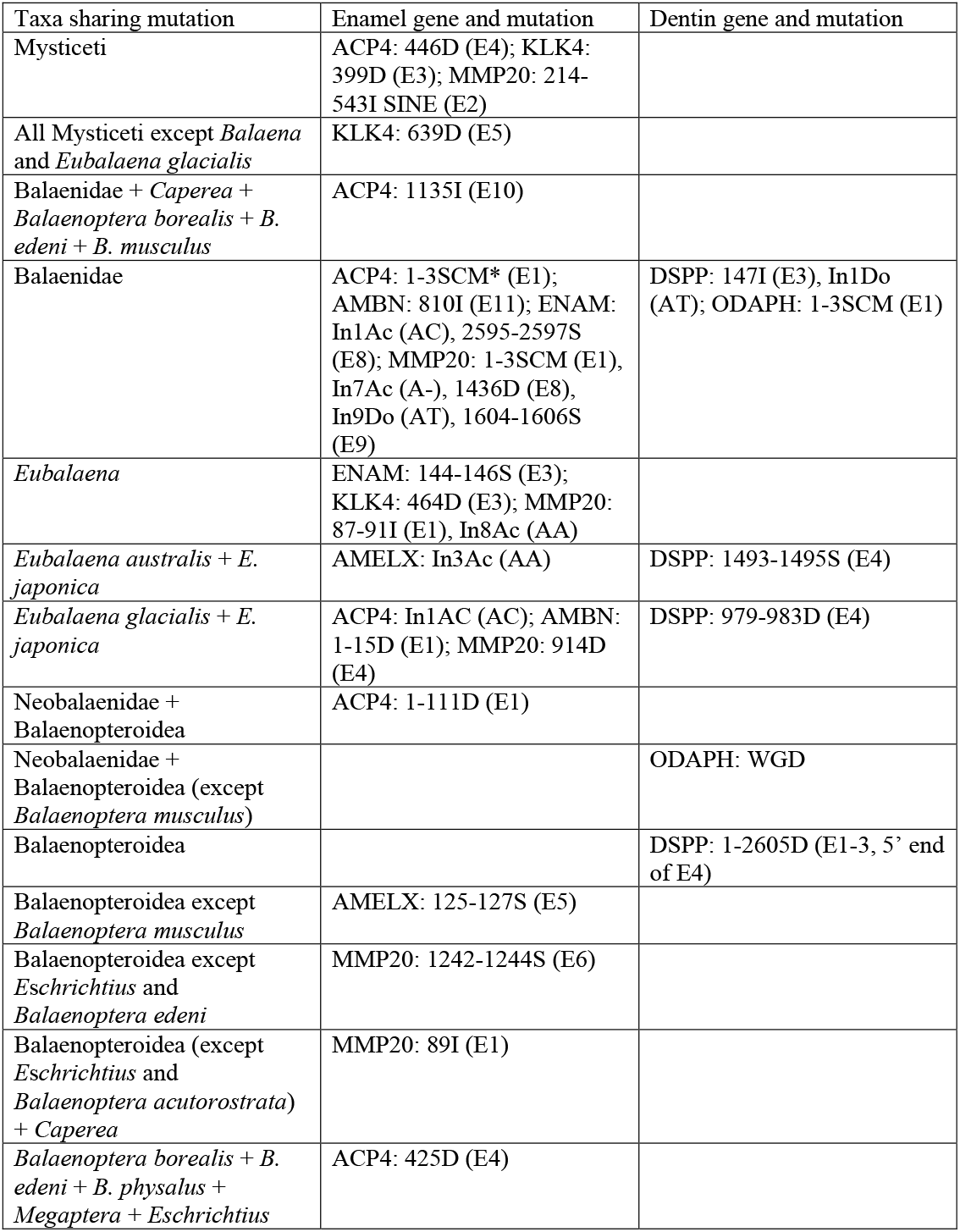
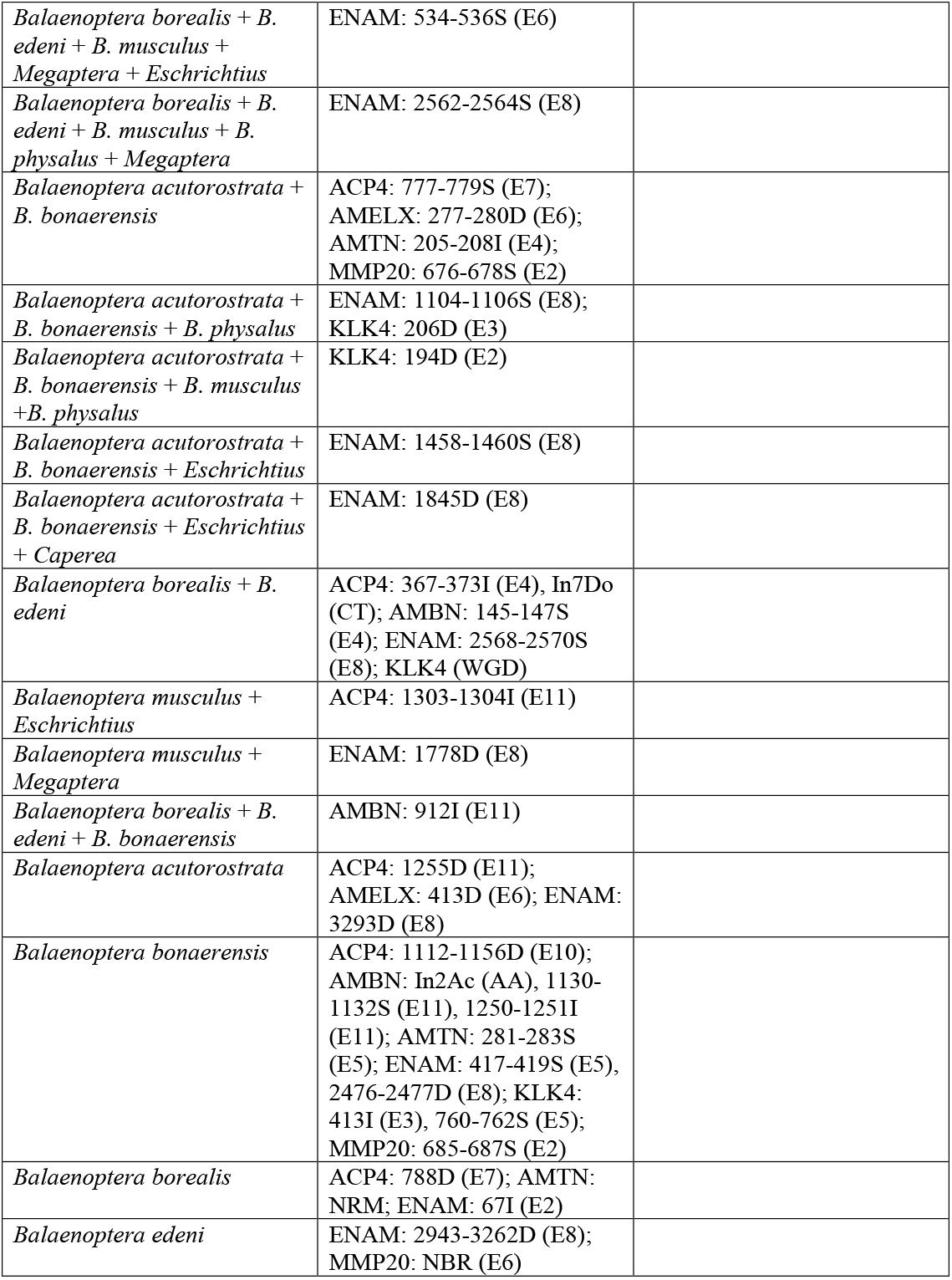
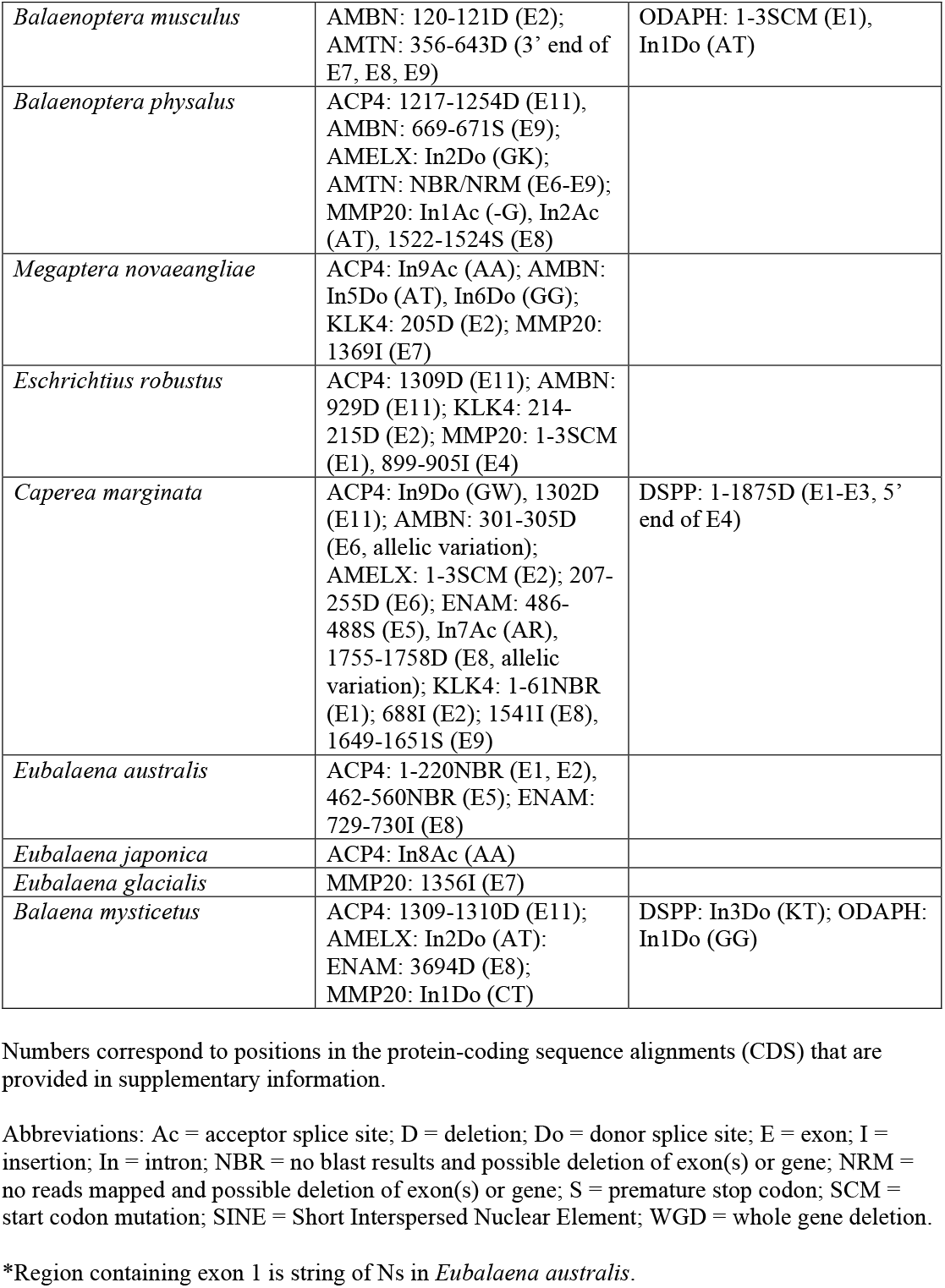
Inactivating mutations in enamel and dentin genes in Mysticeti.

**Fig. 1.**
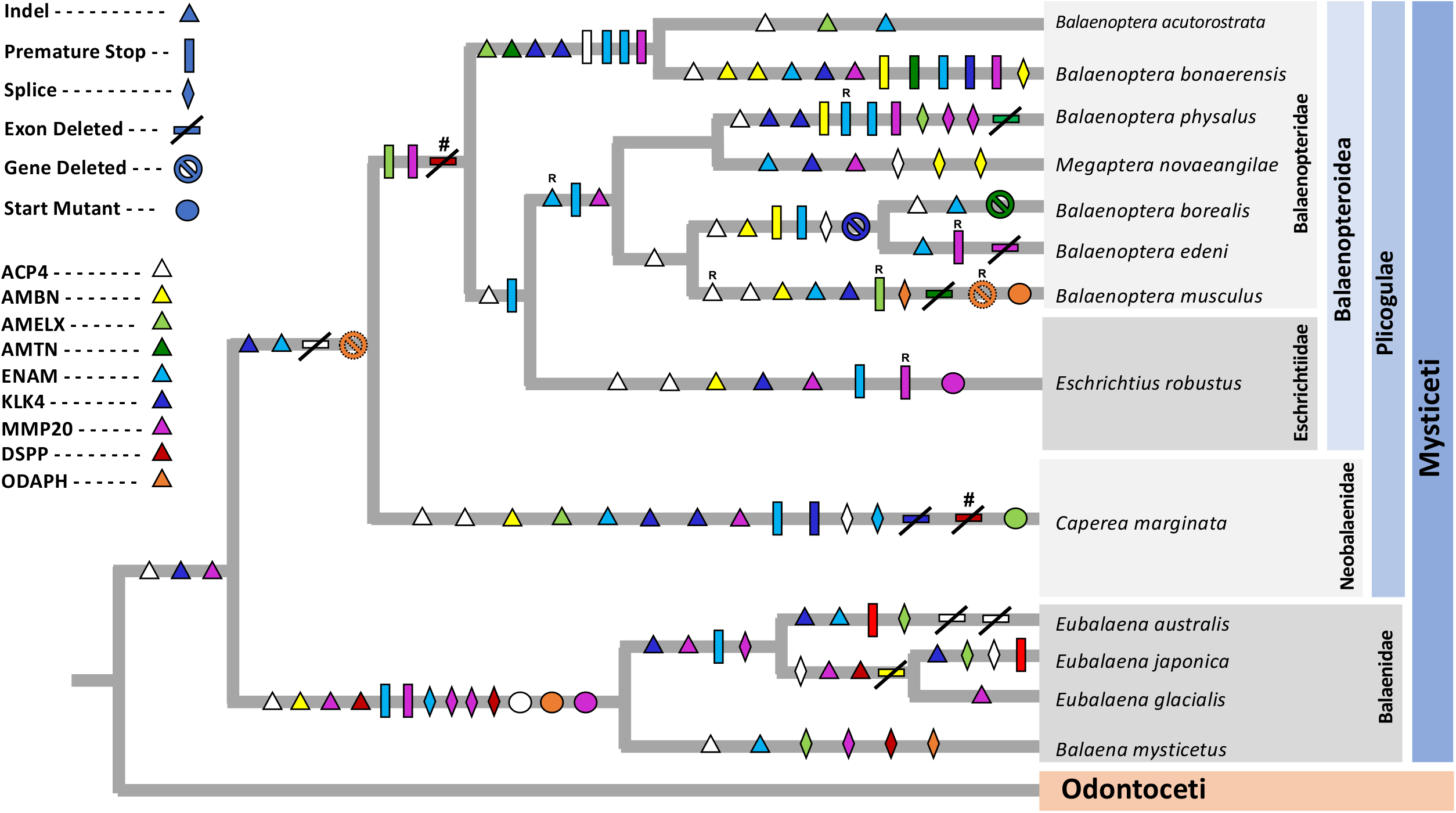
Mapping of inactivating mutations in tooth genes onto a species tree for mysticetes from McGowen et al. (2020a). Inactivating mutations were optimized with DELTRAN. Reversals of inactivating mutations are indicated by an uppercase R above a symbol. In some cases, reversals may be due to lineage sorting of ancestral polymorphism (Springer et al., 2016a) or gene flow among lineages (Árnason et al., 2018). Exons 1-3 and part of exon 4 of *DSPP* are deleted in Plicogulae. This deletion is marked as convergent by pound signs (#) on the stem Balaenopteroidea and *Caperea marginata* branches, but a single deletion that originated in the common ancestor of Plicogulae may instead explain the missing exons (see main text). Abbreviations: Eschr. = Eschrichtiidae, Lipo. = Lipotidae, Mono. = Monodontidae, Neob. = Neobalaenidae, Phoco. = Phocoenidae, Phys. = Physeteridae, Ziph. = Ziphiidae. commenced early history of crown Physeteroidea.

Among the other enamel-related genes, *ENAM* has inactivating mutations in all mysticetes that were examined, but none that are shared by all taxa (Fig. 1). The most inclusive mutations in *ENAM* include a premature stop codon in exon 6 that is shared by *Eschrichtius, Megaptera*, and three species of *Balaenoptera*; a premature stop codon in exon 8 that is shared by *Caperea, Megaptera*, and four species of *Balaenoptera*; and a premature stop codon in exon 8 that is shared by all four balaenids. *AMBN* has one or more inactivating mutations in all mysticetes except for *Balaenoptera acutorostrata* including a 1-bp insertion in exon 11 that is shared by all four balaenids. *AMELX* has one or more inactivating mutations in all mysticetes except for *Balaenoptera musculus* and *Eubalaena glacialis*. The most inclusive inactivating mutation in *AMELX* is a premature stop codon in exon 6 that is shared by all balaenopteroids except for *Balaenoptera musculus*. Finally, *AMTN* remains intact in most mysticetes, but has inactivating mutations in three species of *Balaenoptera* (*B. acutorostrata, B. bonaerensis, B. musculus*) and negative BLAST or map to reference results for the entire protein-coding sequence (*B. borealis*) or exons 6-9 of this gene (*B. physalus*). The only definitive shared inactivating mutation in *AMTN* is a 4-bp insertion in exon 4 in the two minke whales (*Balaenoptera acutorostrata, B. bonaerensis*).

Mu et al. (2021) reported that seven of nine mysticetes that were examined have exon 1 sequences for *ACP4*, including five species with a mutated start codons (GTG). We recovered exon 1 sequences for *ACP4* in three balaenids, but not in any balaenopteroids. The exon 1 sequences that we found are not orthologous with the putative exon 1 sequences that were reported by Mu et al. (2021) for mysticetes. Instead, our analyses indicate that Mu et al.’s (2021) exon 1 sequences for mysticetes are more than 4 kb upstream from the correct exon 1 location in cetaceans and are the result of an annotation error by NCBI’s automated software (Fig. 2A). A ∼2.25 kb deletion in balaenopteroids removed approximately 2.1 kb of sequence that is upstream of exon 1, all of exon 1, and 43 bp at the 5’ end of intron 1 (Fig. 2A). The absence of map to reference results suggests that this exon was also deleted in the neobalaenid *Caperea marginata*.

**Fig. 2.**
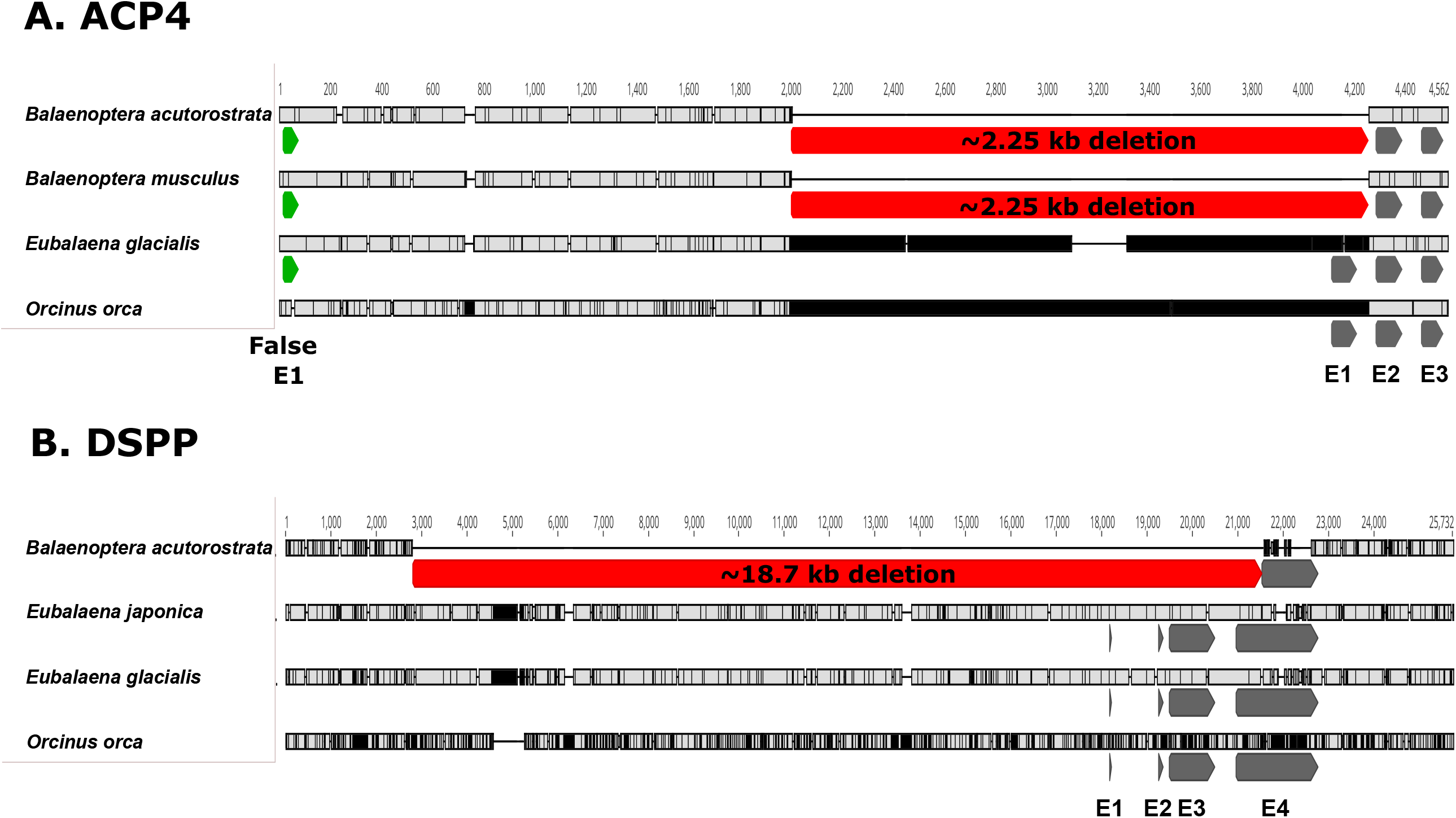
Examples of exon deletions in mysticete tooth genes. **A.** Deletion of exon 1 (E1) of the *ACP4* gene in Balaenopteroidea. The deletion (in red) spans 2.25 kb and includes ∼2.1 kb of sequence that is upstream to exon 1, all of exon 1, and 43-bp at the 5’ end of intron 1. Exons are shown in gray. Mu et al. (2021) reported an incorrect (false) version of exon 1 (in green) in several mysticetes based on misannotations in GenBank. **B.** Deletion of exons 1-3 and part of exon 4 of *DSPP* in a representative balaenopteroid (*Balaenoptera musculus* [blue whale]). Exons 1-3 and part of exon 4 are also deleted in the neobalaenid *Caperea marginata* (not shown). The deletion spans ∼18.7 kb in *B. musculus*.

The tooth/dentin genes *DSPP* and *ODAPH* do not have any inactivating mutations that are shared by all mysticetes. However, *DSPP* exhibits two inactivating mutations that are shared by all four balaenids (1-bp insertion in exon 3, donor splice site mutation [AT] in intron 1) (Table 2). There is also a large deletion (exons 1-3 and most of exon 4) that is present in all species of Plicogulae (Fig. 2B). A caveat regarding the large deletion is that its 3’ boundary is sensitive to alignment settings and may or may not be at a homologous position in balaenopteroids and the neobalaenid *Caperea*. Furthermore, and importantly, it is possible that the 1-bp frameshift insertion in exon 3 of balaenids may have originated in the common ancestor of Mysticeti with subsequent erasure of this evidence for early inactivation on the stem mysticete branch when exon 3 was deleted in balaenopteroids and the neobalaenid. In the case of *ODAPH*, there is a start codon mutation (ATG > TTG) in all four balaenids, complete deletion of the gene in *Caperea marginata* and all balaenopteroids except for *Balaenoptera musculus*, and two inactivating mutations in *B. musculus ODAPH*. The inactivating mutations in dentin/tooth genes suggest that teeth were lost at most two times in crown Mysticeti, i.e., on the stem branches leading to Balaenidae and Plicogulae, respectively (Fig. 1).

### 3.3. Inactivating mutations in odontocetes

All of the enamel genes except for *MMP20* exhibit an inactivating mutation(s) in one or more odontocetes (Table 3, Fig. 3). With the exception of a donor splice site mutation in intron 4 of *Orcinus orca* (killer whale), all of the inactivating mutations are in odontocetes that lack complex enamel with well-developed prisms (*sensu* Werth et al., 2020) and whose enamel (if present) is often worn away in adults. Non-delphinid odontocetes with inactivating mutations in enamel genes include two monodontids (*Monodon monoceros, Delphinapterus leucas*), two phocoenids (*Neophocaena asiaeorientalis, Phocoena phocoena*), one ziphiid (*Mesoplodon bidens*), and three physeteroids (*Physeter macrocephalus, Kogia breviceps, K. sima*). The two monodontids share a donor acceptor splice site mutation (GT -> AT) in intron 2 of *AMELX*. One or both of the monodontids also exhibit autapomorphic inactivating mutations in *ACP4, AMBN, AMELX, AMTN*, and *KLK4*. The two phocoenids share a premature stop codon in exon 2 of *KLK4* and an acceptor splice site mutation (AG -> AT) in intron 2 of *AMTN*. There is also an autapomorphic premature stop codon in *Neophocaena asiaeorientalis AMELX*. Among physeteroids, *Physeter macrocephalus* has three autapomorphic frameshift indels in *ACP4*. The two species of *Kogia*, in turn, exhibit inactivating mutations and/or negative BLAST/map to reference results for five (*K. sima*) or six (*K. breviceps*) enamel genes. Shared inactivating mutations are present in *ACP4, AMELX*, and *ENAM*, and there were negative BLAST results (*K. breviceps*) or no mapped reads (*K. sima*) for *KLK4*. Presumably, *KLK4* is deleted in *Kogia*. Finally, the ziphiid *M. bidens* has three inactivating mutations in *ACP4*.

**Table 3.** Inactivating mutations in enamel genes in Odontoceti.

| Taxa sharing mutation | Enamel gene and mutation |
| --- | --- |
| *Orcinus orca* | ENAM: In4Do (AT) |
| Phocoenidae | KLK4: 73-75S (E2); AMTN: In2Ac (AT) |
| *Neophocaena asiaeorientalis* | AMELX: 386-388S (E6) |
| Monodontidae | AMELX: In2Do (AT) |
| *Monodon monoceros* | ACP4: In6Do (AT); AMBN: 1214-1216S (E11); KLK4: 503-505S (E4) |
| *Delphinapterus leucas* | ACP4: 673D (E7); AMTN: 571D (E8) |
| *Physeter macrocephalus* | ACP4: 425D (E4), 1015D (E9), 1134D (E10) |
| *Kogia* | ACP4: E1-3 (deletion), E11 (deletion); AMELX: 47I (E2), In2Do (GG); ENAM: 2042D (E8), 3429-3430D (E8); KLK4: NBR/NMR (gene deletion) |
| *Kogia breviceps* | ACP4: 657-670D (E7), 901D (E9), 1098-1100S (E10), 1109I (E10); AMBN: In7Ac (AT), In9AC (AT); AMELX: 1-3SCM (E2); AMTN: 373D (E8); ENAM: In6Do (CT); |
| *Kogia sima* | AMBN: NRM (gene deletion); ENAM: NRM (E1-E6); |
| *Mesoplodon bidens* | ACP4: 146-149I (E2), 536-543D (E5), In8Ac (GG) |

Numbers correspond to positions in the protein-coding sequence alignments (CDS) that are provided in supplementary information.
Abbreviations: Ac = acceptor splice site; D = deletion; Do = donor splice site; E = exon; I = insertion; In = intron; NBR = no blast results and possible deletion of exon(s) or gene; NRM = no reads mapped and possible deletion of exon(s) or gene; S = premature stop codon; SCM = start codon mutation.

**Fig. 3.**
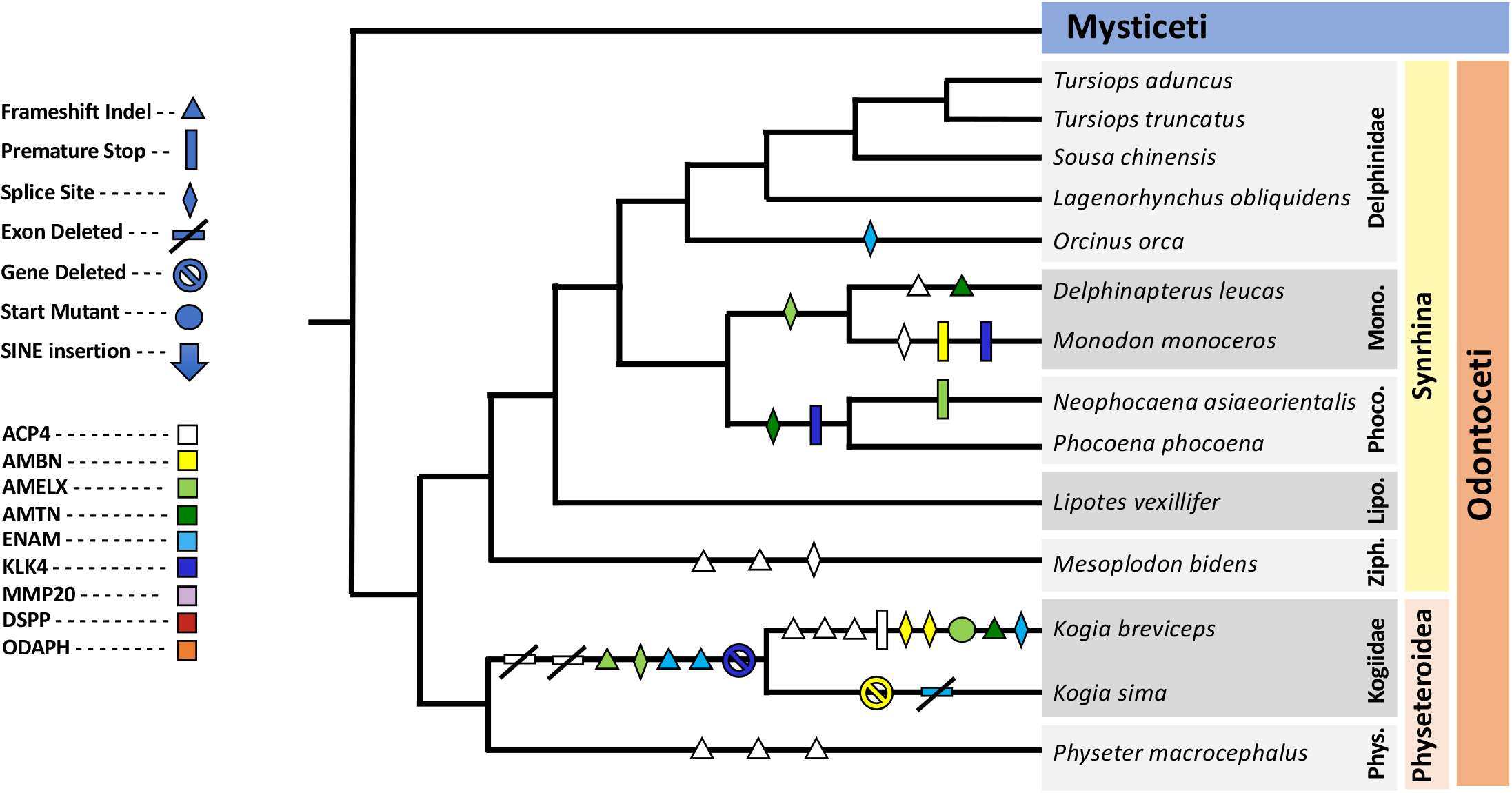
Mapping of inactivating mutations in tooth genes onto a species tree for odontocetes from McGowen et al. (2020a). Inactivating mutations were optimized with DELTRAN.

### 3.4. Selection analyses

Selection analyses were performed on seven enamel genes that were concatenated together and two tooth/dentin genes that were concatenated together (Supplementary Information). The concatenated alignment of enamel genes included 44 taxa and 9618 bp. The concatenated dentin alignment included 35 taxa and 2127 bp. Balaenopteroids and Plicogulae were excluded from the dentin alignment because of their incompleteness. The results of selection analyses on both the enamel and dentin genes are summarized in Table 4.

**Table 4.**
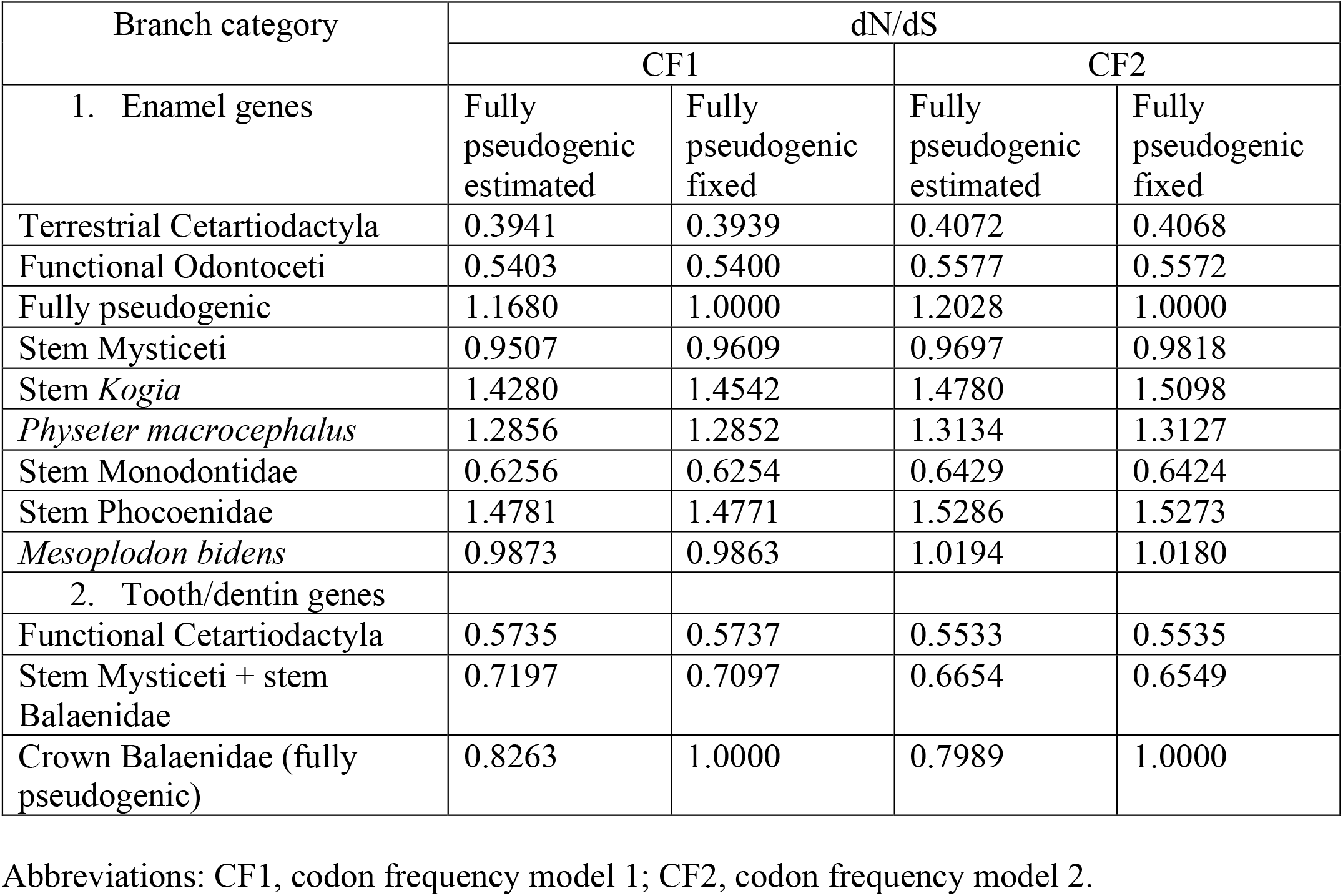
Selection analyses on seven enamel genes and two tooth/dentin genes.

For the enamel genes, dN/dS values for the two background categories, terrestrial cetartiodactyls and odontocetes with complex enamel (Werth et al., 2020), ranged from 0.3939 to 0.4072 for the former and 0.5400 to 0.5577 for the latter. These values are indicative of purifying selection and further suggest that purifying selection has been stronger in terrestrial cetartiodactyls than in odontocetes with complex enamel, perhaps because odontocetes generally do not masticate their food as assiduously as herbivorous/omnivorous terrrestrial cetartiodactyls. The dN/dS value for fully pseudogenic branches ranged from 1.168 (CF1) to 1.2028 (CF2) when these values were estimated rather than fixed at 1.0. These values are significantly different from 1.0 based on log likelihood ratio tests (p = 0.029 [CF1], p = 0.009 [CF2]). Among the transitional branches, the dN/dS value on the stem mysticete branch was intermediate between the functional and pseudogenic dN/dS values and ranged from 0.9507 to 0.9818. DN/dS values for transitional branches in Odontoceti were all higher than dN/dS values for the two functional categories and in three cases (stem *Kogia, Physeter*, stem Phocoenidae) were slightly higher than the estimated dN/dS values for the fully pseudogenic branches. However, in each case these elevated values were not significantly higher than the estimated values for the pseudogenic branches when individual transitional branches were constrained to have the same dN/dS value as the fully pseudogenic branches (stem *Kogia*: p = 0.48 [CF1], p = 0.47 [CF2]; *Physeter*: p = 0.74 [CF1], p = 0.76 [CF2]; stem Phocoenidae: p = 0.42 [CF1], p = 0.41 [CF2]).

Selection analyses on the two dentin genes yielded dN/dS values of 0.5735 (CF1) and 0.5533 (CF2) for the background branches that lead to dentate taxa, 0.8263 (CF1) and 0.7989 (CF2) for the fully pseudogenic branch category of crown mysticetes that are edentulous, and 0.7179 (CF1) and 0.6654 (CF2) for the transitional branch that merges the stem Mysticeti and stem Balaenidae branches (Table 4). As expected, the background branches have the lowest dN/dS values, the fully pseudogenic branches have the highest dN/dS values, and the transitional branch has intermediate dN/dS values. The dN/dS value for the fully pseudogenic crown branches is less than the expected value of 1.0, but the difference is not statistically significant based on a likelihood ratio test (p = 0.54 [CF1], p = 0.47 [CF2]).

### 3.5. Gene inactivation times

The timings of enamel and dentin/tooth loss were estimated by proxy using dN/dS values for the concatenations of enamel and dentin/tooth genes, respectively, equations from Meredith et al. (2009), and cetacean divergence dates from McGowen et al. (2020a). The mean of eight different estimates for inactivation of the enamel-specific genes on the stem Mysticeti branch is 34.62 Ma, whereas the mean inactivation time for dentin/tooth-specific genes on the combined stem Mysticeti + stem Balaenidae branch is 19.94 Ma (Table 5, Fig. 4). Note that species in Plicogulae were not included in the dentin/tooth gene analysis because of exon and gene deletions. These inactivation times suggest that enamel was lost very early on the stem Mysticeti branch, whereas dentin/teeth were lost on the stem Balaenidae branch, which extends from 25.73 Ma to 10.61 Ma based on McGowen et al.’s (2020a) autocorrelated rates timetree (Fig. 4). This is consistent with the observation that shared inactivating mutations occur in Balaenidae but not for all mysticete species. There is insufficient information to estimate a reliable date for dentin/tooth loss in the ancestor of Plicogulae based on dN/dS analyses, but exon deletions in *DSPP* and the complete deletion of *ODAPH* in most plicogulans suggest that dentin/teeth were lost on the stem plicogulan branch, which extends from 25.73 to 22.11 million years ago based on McGowen et al.’s (2020a) timetree. Overall, these results suggest a two-step model for the loss of teeth in the ancestry of living baleen whales, where the initial loss of enamel on the mysticete stem lineage was followed by subsequent loss of dentin/teeth on two or more lineages within the crown group.

**Table 5.** Inactivation times (Ma) for seven enamel genes on the stem Mysticeti branch and two dentin/tooth genes on the stem Mysticeti + stem Balaenidae branches

| Gene concatenation | CF1 |  |  |  | CF2 |  |  |  | Mean Inactivation Time |
| --- | --- | --- | --- | --- | --- | --- | --- | --- | --- |
|  | dN/dS estimated |  | dN/dS = 1 |  | dN/dS estimated |  | dN/dS = 1 |  |  |
|  | 1 syn rate | 2 syn rates | 1 syn rate | 2 syn rates | 1 syn rate | 2 syn rates | 1 syn rate | 2 syn rates |  |
| Seven enamel genes | 33.63 | 32.79 | 36.01 | 35.73 | 33.50 | 32.63 | 36.38 | 36.24 | 34.62 |
| Two dentin genes | 25.71 | 23.40 | 18.94 | 17.06 | 22.53 | 20.28 | 16.54 | 15.06 | 19.94 |

Abbreviations: CF1, codon frequency model 1; CF2, codon frequency model 2; dN = nonsynonymous substitutions per nonsynonymous site; dS = synonymous substitutions per synonymous site; Ma = millions of years ago, syn = synonymous.

**Fig. 4.**
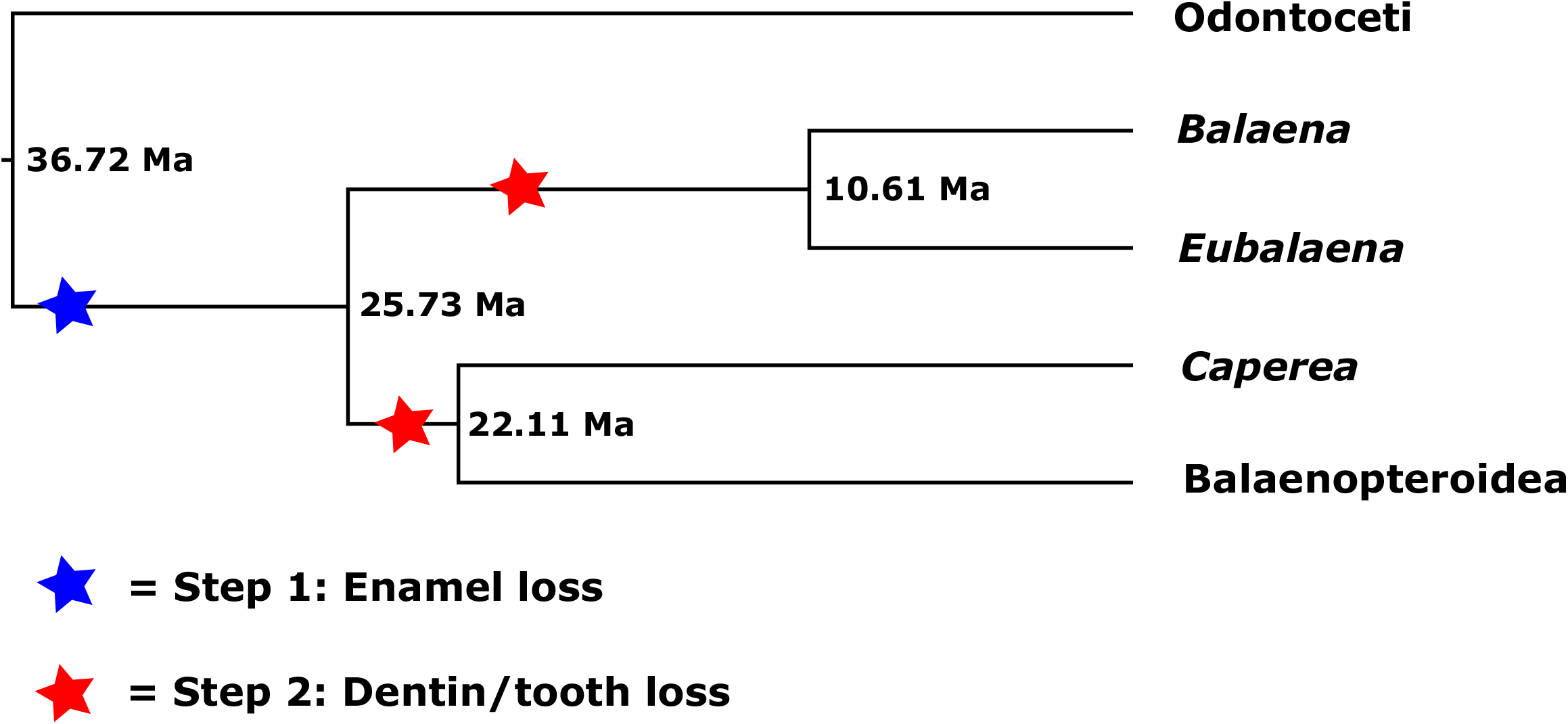
Two-step model for tooth loss in extant mysticetes based on shared inactivating mutations and gene inactivation dating. **Step 1:** Enamel was lost from the teeth early on the stem mysticete branch (blue star). Shared inactivating mutations in three genes (*ACP4, KLK4, MMP20*) provide support for enamel loss on the stem mysticete branch, and gene inactivation dating based on seven enamel-related genes places enamel loss at ∼35 million years ago. **Step 2:** Dentin and teeth were lost independently on the stem Balaenidae and stem Neobalaenidae + Balaenopteroidea branches, respectively (red stars). Shared inactivating mutations provide support for dentin/tooth loss on these two branches, and gene inactivation dating based on two dentin/tooth-related genes places the loss of teeth at ∼20 MYA on the stem Balaenidae branch. Gene inactivation dating was not applied to the stem Balaenopteroidea + Neobalaenidae branch because exons 1-3 of DSPP are deleted in all plicogulans. *Caperea* belongs to Neobalaenidae; Balaena and *Eubalaena* belong to Balaenidae. Divergence dates at nodes are from McGowen et al. (2020a).

## 4. Discussion

### 4.1. Tooth loss in mysticetes

Both fossil evidence and ancestral reconstructions of edentulism suggest that postnatal teeth were lost on the stem lineage to crown Mysticeti. Previous studies based on molecular data validate the hypothesis that the genetic toolkit for enamel production was knocked out on the stem mysticete branch (Meredith et al., 2011; Gatesy et al., 2021; Mu et al., 2021). However, shared inactivating mutations in a dentin-related gene have not been reported in mysticetes. Indeed, it remains unclear if enamel and dentin loss were coupled or if enamel loss preceded edentulism. Gatesy et al. (2021) articulated three hypotheses for the evolution of edentulism in extant baleen whales: (H1) enamel and dentin components of teeth were lost simultaneously on the stem mysticete branch, (H2) enamel and dentin/teeth were lost in a stepwise fashion on the stem mysticete branch (i.e., enamel first, then dentin/teeth), and (H3) enamel loss occurred on the stem mysticete branch followed by the independent loss of dentin (and teeth) in two or more crown mysticete lineages. We addressed these hypotheses through molecular evolutionary analyses of protein-coding sequences for seven enamel-related genes and two tooth/dentin-related genes.

In the case of the seven enamel-related genes we inferred three inactivating mutations that are shared by all mysticetes: a SINE insertion in exon 2 of *MMP20*, a 1-bp deletion in exon 4 of *ACP4*, and a 1-bp deletion in exon 3 of *KLK4*. Each of these inactivating mutations has previously been reported albeit with less comprehensive taxon sampling for mysticetes (*MMP20* [Meredith et al., 2011]; *ACP4* [Mu et al., 2021]; *KLK4* [Gatesy et al., 2021]). In addition to these inactivating mutations that are shared by all mysticetes, there are numerous mutations, some of which are shared by two or more species, in all seven of the enamel-related genes (Fig. 1; Table 2). These results suggest that enamel-related genes have been evolving neutrally since the loss of enamel on the stem mysticete branch. The absence of shared inactivating mutations (for Mysticeti) in four enamel-related genes (*AMBN, AMELX, AMTN, ENAM*) may reflect a lag time between the onset of neutral evolution and the accumulation of the first inactivating mutation in each of these four genes. This scenario has previously been inferred for cone-specific phototransduction genes in rod monochromatic cetaceans (Springer et al., 2016b). DN/dS analyses validate the hypothesis that the seven-enamel related genes have evolved neutrally in crown Mysticeti after purifying selection was abolished on the stem mysticete branch.

By contrast with the seven enamel-related genes, we did not find inactivating mutations that are shared by all mysticetes in either of the dentin/tooth related genes (*DSPP, ODAPH*) that we examined. Springer et al. (2016a) previously reported this result for *ODAPH* (= *C4orf26*) but with more limited taxon sampling. Nevertheless, there are inactivating mutations in *ODAPH* that map to the common ancestors of Balaenidae (start codon mutation) and Plicogulae (gene deletion), respectively, although in the latter case one balaenopterid (*Balaenoptera musculus*) retains a pseudogenic copy of *ODAPH*. One explanation for this pattern is that the deletion of *ODAPH* was polymorphic in the ancestor of Plicogulae with subsequent lineage sorting (Springer et al., 2016a), but alternatively, the presence of *ODAPH* in *B. musculus* might be due to gene flow and introgression, which is thought to have been extensive in mysticete phylogeny (Berube and Aguilar, 1998; Arnason et al., 2018; Springer et al., 2020). In the case of *DSPP*, a splice site mutation in intron 1 and a 1-bp insertion in exon 3 are shared by all four balaenids. There is also a large deletion that is shared by Plicogulae. This deletion encompasses exons 1-3 and most of exon 4. The most parsimonious hypothesis is that this deletion occurred in the common ancestor of Plicogulae. However, alignment uncertainties at the 3’ boundary of this deletion, which occurs in a highly repetitive region of *DSPP*, allow for the less likely scenario wherein the deletion occurred independently in Neobalaenidae and in Balaenopteroidea. Either way, the deletion of exon 3 in Plicogulae leaves open the possibility that the 1-bp frameshift insertion in Balaenidae originated in the common ancestor of Mysticeti with subsequent loss of evidence for this mutation in Plicogulae when exon 3 was deleted in this clade. In summary, the combined evidence from *ODAPH* and *DSPP* suggests that the toolkit for tooth/dentin formation was inactivated at most two times within crown Mysticeti, once in the common ancestor of Balaenidae and once in the common ancestor of Plicogulae.

Given the very slow rates of molecular evolution that occur in mysticetes (Meredith et al., 2009), selection on the two dentin/tooth genes may have been relaxed in the common ancestor of Mysticeti even though there are no shared inactivating mutations in tooth/dentin genes in this clade. We tested this hypothesis using coding sequences for *ODAPH* and the first three exons of *DSPP* in balaenids (exon 4 is highly repetitive and difficult to align). The results of these analyses suggest that selection on the two tooth/dentin genes was relaxed ∼20 Ma in the common ancestor of Balaenidae, which is ∼15 million years after selection was relaxed on the seven enamel genes in the common ancestor of crown Mysticeti. Together, shared inactivating mutations and the results of dN/dS analyses provide support for H3 (see above) wherein enamel loss and tooth loss were decoupled in the ancestry of living baleen whales (Fig. 4). First, enamel loss occurred ∼35 Ma on the stem mysticete branch. Next, independent dentin/tooth loss occurred on the stem balaenid and stem Plicogulae branches, respectively. Tooth/dentin loss occurred ∼20 Ma on the stem balaenid branch based on dN/dS analyses and inactivation dating. We were unable to date tooth/dentin loss on the stem Plicogulae branch using our dN/dS methods, but if the loss of *ODAPH* and/or the deletion of most of *DSPP* are shared inactivating mutations in Neobalaenidae + Balaenopteroidea, then dentin/teeth may have been lost on this branch. Estimated divergence dates from McGowen et al.’s (2020a) timetree for Cetacea suggest that dentin/tooth loss on the stem Plicogulae branch occurred 25.73-22.11 Ma.

Our hypothesis for tooth loss (H3) in the ancestry of living whales is incongruent with ancestral reconstructions of edentulism in mysticetes (Fitzgerald, 2006, 2010; Deméré et al., 2008; Meredith et al., 2011a). One explanation for this incongruence is the possibility that small enamelless teeth were set in the gums rather than in bony alveoli in some stem mysticetes and crown mysticetes. This condition is known in some extant ziphiids (Boschma, 1951; Fordyce et al., 1979; Gomerčić et al., 2006). In addition, Lambert et al. (2008) suggested that small teeth may have been embedded in the gums rather than in alveoli in the Miocene ziphiid *Nazcacetus urbinai*. If this condition also occurred in late stem and early crown mysticetes from the Oligocene and Miocene, it is possible that fossils might not record this anatomical condition. The small, loosely set teeth may have become detached from the jaws of such species by post-mortem taphonomic processes and not recovered in association with the skull (Gatesy et al., 2021). Along these lines, there is suggestive but inconclusive evidence for the presence of very small teeth, set in alveoli, that were located at the tips of the upper and lower jaws in some Oligocene eomysticetid mysticetes including *Yamatocetus canaliculatus* and *Tokarahia* sp. (Okazaki, 2012; Boessenecker and Fordyce, 2015). It remains to be determined if this condition may have occurred in some of the earliest crown mysticetes. More generally, additional paleontological and molecular evolutionary studies will be required to test competing hypotheses pertaining to the loss of teeth in extant mysticetes. Beyond *DSPP* and *ODAPH* we are unaware of additional genes that are dentin/tooth-specific but not enamel-specific. The discovery of such genes, if they exist, will allow for a more thorough investigation of the timing and pattern of tooth loss in mysticetes.

Molecular evolutionary evidence for asynchronous enamel and tooth loss in mysticetes, with the latter occurring within the crown group, is compatible with the co-occurrence hypothesis for the evolution of baleen wherein there were transitional forms such as *Aetiocetus weltoni* that possessed both teeth and baleen (Deméré et al., 2008; Ekdale and Deméré, 2021). More specifically, the evolution of baleen occurred on the stem mysticete branch before teeth were finally lost within crown Mysticeti (Fig. 4). By contrast, our results are incompatible with the toothless suction-feeding hypothesis wherein teeth were lost prior to the evolution of baleen (Peredo et al., 2017, 2018). This second hypothesis requires the loss of teeth very deep in the mysticete tree on the stem lineage, while our analysis of tooth genes implies a much later loss of teeth within crown Mysticeti. Peredo et al. (2018) argued that the Oligocene mysticete *Maiabalaena nesbittae* (∼33 Ma) exemplified the toothless, baleenless, suction-feeding intermediate that bridged the transition from feeding with teeth on single prey items to batch filter-feeding with baleen. However, character optimizations based on the mysticete data matrix compiled by Peredo et al. (2018) imply that *Maiabalaena* does not represent the ancestral condition and instead is more likely an aberrant side branch that may have independently lost the postnatal dentition (Gatesy et al., 2021) millions of years prior to convergent tooth loss in crown Mysticeti (Fig. 4).

Thewissen et al. (2017) reported the presence of a mineralized dentin matrix, but not an enamel matrix, in tooth germs of fetal specimens of *Balaena mysticetus* (bowhead) that reach the bell stage of tooth development before they are resorbed. Transitory tooth buds that reach the bell stage have also been reported for *Balaenoptera physalus* (fin whale) (Dissel-Scherft and Vervoort, 1954; Deméré et al., 2008) and *Balaenoptera acutorostrata* (Antarctic minke whale) (Ishikawa and Amasaki, 1995). In the case of *B. acutorostrata*, Ishikawa and Amasaki (1995) reported the presence of predentin that appears in the bell stage of tooth development. Ishikawa and Amasaki (1995) also noted that both the inner and outer enamel layers form during the bell stage, although the inner enamel layer never differentiates into ameloblasts. Instead, the tooth buds begin to degenerate between dentin formation and ameloblast differentiation. Our results suggest that the predentin/dentin of both Ishikawa and Amasaki (1995) and Thewissen et al. (2017) is a degenerative form of dentin because of inactivating mutations that are present in mysticete *DSPP*. Indeed, type 1 collagen and DSPP are the most abundant protein components of dentin (Yamakoshi, 2009) and of these only DSPP is dentin specific. It will be interesting to learn if any remnants of *DSPP* are expressed in the dentin matrix of fetal mysticetes. The most likely candidate would be a remnant of dentin phosphoprotein (DPP), which is encoded by the 3’ region of exon 4 and is one of the three main components of dentin sialophosphoprotein (DSPP) along with dentin sialoprotein (DSP) and dentin glycoprotein (DGP) (Yamakoshi, 2009; Yamakoshi and Simmer, 2018). [DPP, DSP, and DGP are the result of protease-processed cleavage of the full length DSPP protein.] It also remains to be determined if the tooth buds of fetal whales have a role in guiding the formation of the baleen racks (Thewissen et al., 2017) or if they are merely rudiments of the formerly functional dentition that currently serve no purpose.

### 4.2. Enamel loss in odontocetes

Odontocetes exhibit a wide range of enamel phenotypes including highly prismatic enamel with or without HSB, intermediate enamel with less distinct prisms and amorphous crystallite aggregations that run in multiple directions, thin prismless enamel that is easily removed due to wear, and the complete absence of enamel (Ishiyama, 1987; Bianucci and Landini, 2006; Meredith et al., 2009, 2013; Loch et al., 2013a,b; Werth et al., 2020). There is even a recently discovered toothless extinct species, the dwarf dolphin *Inermorostrum xenops*, which belongs to the early diverging odontocete clade Xenorophidae (Boessenecker et al., 2017).

Living odontocetes with highly prismatic enamel (with or without HSB) include delphinids and river dolphins (Werth et al., 2020). These taxa use their teeth to procure, retain, and process their prey more than do other odontocetes (Werth et al., 2020). Our study included five delphinids and one river dolphin (*Lipotes vexillifer*). We are not aware of enamel microstructure data for *L. vexillifer*, but this taxon has highly crenulated enamel (Brownell and Herald, 1972) as in the closely related Amazon River dolphin (*Inia geoffrensis*). In *Inia*, the enamel is highly prismatic with HSB (Werth et al., 2020). Among delphinids and river dolphins in our study, the only inactivating mutation in the seven enamel-related genes is a donor splice site mutation (GT to AT) in intron 4 of *ENAM* in *Orcinus orca*. The other eight odontocetes that were included in our study include two phocoenids, two monodontids, one ziphiid, and three physeteroids. Of these, the phocoenids have intermediate enamel and exhibit inactivating mutations in one (*Neophocaena asiaeorientalis*) to three (*Phocoena phocoena*) of the enamel genes. With the exception of *Tasmacetus shepherdi* (Shepherd’s beaked whale), which has a full set of teeth, the dentition in extant ziphiids is restricted to tusks and occasionally small vestigial teeth (Loch and van Vuuren, 2016). Four ziphiids that have been investigated (*Berardius bairdii, B. arnuxii, Mesoplodon densirostris, Ziphius cavirostris*) have no enamel or only a thin layer of enamel on their tusks (Ishiyama, 1987; Loch and van Vuuren, 2016; Thewissen, 2018; Werth et al., 2020). The single ziphiid that was included in our study, *Mesoplodon bidens*, exhibits multiple inactivating mutations but all of these are confined to one (*ACP4*) of the seven enamel genes. Among the monodontids, *Monodon monoceros* lacks enamel and *Delphinapterus leucas* has prismless enamel (Ishiyama, 1987). *M. monoceros* has inactivating mutations in four enamel genes and *D. leucas* has inactivating mutations in three enamel genes. The presence of a shared inactivating mutation in *AMELX* in these taxa suggests that enamel degeneration commenced in their common ancestor.

In the case of physeteroids, Bianucci and Landini (2006) reported the presence of enamel in several stem taxa (*Zygophyseter, Naganocetus, Aulophyseter*) and suggested that enamel was lost in the last common ancestor of crown Physeteroidea, which includes two extant genera (*Kogia, Physeter*) as well as several extinct forms (*Orycterocetus, Placoziphius, Physeterula, Scaphokogia*) that were coded by these authors as having no enamel. Other studies have also reported the absence of enamel in *Physeter* (Flower and Lydekker, 1891) and *Kogia* (Willis and Baird, 1998). By contrast, some authors have noted the presence of a thin layer of prismless enamel in both *Physeter* (Ishiyama, 1987) and *Kogia* (Plön, 2004; Bloodworth and Odell, 2008; Werth et al., 2020). The apparent discrepancy between different studies may be explained by the localized occurrence of very thin enamel at the tips of the teeth and the rapid erosion of this enamel once the teeth have erupted (Ishiyama, 1987; Plön, 2004; Bloodworth and Odell, 2008). At the molecular level, all three extant physeteroid species have inactivating mutations in at least one of their enamel genes. All of the inactivating mutations in *P. macrocephalus* are in *ACP4*, as was also observed for *M. bidens*. For *Kogia*, six of the seven enamel genes have inactivating mutations or have been completely deleted in one or both species. However, there are no mutations that are shared by *Physeter* and *Kogia*. Nevertheless, the relatively high dN/dS values on the *Physeter* and stem *Kogia* branches (Table 4) are consistent with the hypothesis that enamel degeneration from prismatic enamel to prismless enamel occurred very early in the history of crown Physeteroidea.

In summary, nine of 14 odontocete species that were included in our study have one or more inactivating mutations in their battery of seven enamel genes. Odontocete species with absent or degenerative enamel exhibit many more inactivating mutations than odontocetes with highly prismatic enamel such as delphinids, most of which have intact protein-coding sequences and splice sites for all seven enamel genes. These results also demonstrate that thin, prismless enamel can still be manufactured with a defective genetic toolkit for enamel production.

## Acknowledgments

This work was supported by National Science Foundation grant DEB-1457735 (J.G., M.S.S.). Paintings of animals are by C. Buell. We thank the Southwest Fisheries Science Center (SWFSC) and the National Marine Mammal and Sea Turtle Research (MMASTR) Collection for providing *Caperea marginata* DNA (SWFSC Lab ID 5989). We also thank the SWFSC, the National Marine Fisheries Service, and the Pacific Islands Fisheries Science Center (Erin Oleson, Chief Scientist) for providing *Kogia sima* DNA (SWFSC Lab ID 175303).

